# Magnitude and mechanism of siderophore-mediated competition in bacterial interactions

**DOI:** 10.1101/094094

**Authors:** Konstanze T. Schiessl, Elisabeth M.-L. Janssen, Stephan M. Kraemer, Kristopher McNeill, Martin Ackermann

## Abstract

Whether microbial interactions are predominantly cooperative or competitive is a central question in microbial ecology, and determines the composition and stability of microbial communities. The secretion of iron chelators called siderophores is a model system for cooperative interactions, even though these chelators can also mediate competition by depriving competitors of iron. Using a genetically engineered experimental system based on the *Pseudomonas aeruginosa* siderophore pyochelin, we found that secreting siderophores to inhibit a competitor can lead to higher benefits than secreting siderophores to make iron available. Based on thermodynamic modeling, we propose that competitive inhibition by siderophores is efficient in kinetically controlled saturated systems, where dissolution of precipitated iron phases is slow. Under these conditions, met in many natural environments, secreted siderophores temporarily reduce the concentration of available iron and can thus induce growth inhibition in a competing strain, even at high iron concentrations. These findings give insight into the function of siderophores: In addition to its cooperative nature, siderophore secretion could also be a widespread mechanism for mediating competitive interactions. Our functional investigation reveals a complexity in microbial interaction networks that would remain hidden when focusing on genomic information alone.

## Introduction

Apart from rare exceptions, all bacteria rely on iron for survival and growth (Andrews *et al.,* 2003). As a consequence, competition for iron is a key factor in interspecies interactions (Simões *et al.,* 2008; Trejo Hernandez *et al.*, 2014; Traxler *et al.*, 2012). A common bacterial iron uptake strategy relies on secretion of iron chelators called siderophores. The primary role of siderophores is commonly thought to be increasing iron availability, e.g., to overcome low iron solubility in aerobic, pH-neutral environments (Andrews *et al.*, 2003). However, siderophores could also play a role in iron competition, since distinct and specific receptors are necessary for the uptake of different iron-siderophore complexes (Wandersman and Delepelaire, 2004). Secreted siderophores could thus sequester iron that would otherwise be freely available, leading to growth inhibition of strains unable to access the chelated iron. Several mathematical models of the effects of siderophores on competitive interactions support this view (Eberl and Collinson, 2009; Fgaier and Eberl, 2010; 2011). Also, inhibition of bacterial growth has been linked to the presence of siderophores in various natural and engineered environments (Vachée *et al.*, 1997; Gram *et al.*, 1999; Khare and Tavazoie, 2015).

Despite these findings, siderophore secretion has mostly been studied as an example of cooperative behavior (Velicer, 2003; West *et al.*, 2006). Secreted siderophores are available to surrounding cells that express the cognate receptor, e.g., to so-called “cheaters” that do not contribute to siderophore production (Griffin *et al.*, 2004) (Ross Gillespie *et al.*, 2007) (Brockhurst *et al.*, 2008). Based on the use of siderophores as a model system for cooperation in laboratory systems, the distribution of siderophore production and receptor genes in the environment has been interpreted mainly as a result of cooperative dynamics (Cordero *et al.*, 2012; Andersen *et al.*, 2015), although they could also reflect competitive dynamics. Since the nature of interactions between different microorganisms in the environment strongly influences the fate of microbial communities (Coyte *et al.*, 2015; Oliveira *et al.*, 2015), differentiating between these different types of interactions is important.

Here, our goal was to contribute to this understanding by quantifying the competitive effect of secreted siderophores in a controlled laboratory system. We developed an experimental system with genetically engineered *Pseudomonas aeruginosa* strains that only differed in the ability to access iron chelated by the siderophore pyochelin. We found that even a low concentration of a this comparatively weak siderophore can efficiently inhibit growth of a competitor strain. Our data also show that the benefits conveyed by inhibiting growth can surpass the benefits mediated by making iron more available in the absence of a competitor. In addition, using thermodynamic modeling, we propose that the mechanism for growth inhibition is based on kinetics: Siderophores can temporarily deplete dissolved iron, slowing down iron uptake of freely-dissolved iron. This depletion strategy makes siderophores efficient at inhibiting growth of competitors even at high iron concentrations.

## Material and Methods

### Strains

We used a ‘secretor’ strain (pyochelin production and uptake), a ‘recipient’ strain (no siderophore production) and a ‘nonrecipient’ strain (neither siderophore production nor uptake) based on *Pseudomonas aeruginosa* PAO1 (ATCC 15692). More specifically, the secretor was knocked out in a gene necessary for production of the siderophore pyoverdine (PAO1 Δ*pvdD*) and obtained from Pierre Cornelis’ laboratory (Ghysels *et al.*, 2004). Thus, the secretor can only produce one type of siderophore, pyochelin. The recipient was a Δ*pvdD*Δ*pchEF* double-knockout in siderophore production, also from (Ghysels *et al.*, 2004). For the nonrecipient, we further used transduction to delete the *fptA* gene necessary for uptake of pyochelin (see SI). To distinguish the strains in competition, they were tagged with constitutively expressed fluorophores. The GFP and mCherry marker were integrated using the Tn7 system (Lambertsen *et al.*, 2004). The expression was under control of the constitutive promoter P_A1/04/03_, a derivate of the lactose promoter.

### Media and growth conditions

For all growth studies, we used succinate minimal medium (7.58 mM (NH_4_)_2_SO_4_, 0.81 mM MgSO_4_*7H_2_O, 33.87 mM succinic acid, 25 mM HEPES). The pH was adjusted to 7.2, and after autoclaving, 10 mM K_2_HPO_4_ were added from a sterile stock solution. Directly before incubation, the media was filtered with a 0.1-*μ*m filter to reduce the amount of precipitated background iron (Cellulose Nitrate Membranes, 0.1 μm, 47 mm ø, Whatman). FeCl_3_ (anhydrous, 97% reagent grade) and FeCitrate (Iron(III) citrate tribasic monohydrate, 18−20% Fe basis) were added as iron sources. 1000x stock solutions were made in 0.1 M HCl for FeCl_3_ and distilled H_2_O for FeCitrate, filter sterilized, and kept at 4 °C. All chemicals were obtained from Sigma Aldrich. A standard of pyochelin was obtained from EMC microcollections (Tübingen, Germany), dissolved in methanol (HPLC reagent grade) and stored at −20°C. Working stocks were made with sterile nanopure water.

Strains were streaked from a frozen stock on standard LB agar. Single colonies were picked and incubated for 20 hours in succinate minimal medium containing 2 *μ*M FeCitrate at 37 °C in an orbital shaker (220 rpm). From dilutions of these cultures, both monoculture as well as competition experiments were started. For all experiments, starting cell density was adjusted based on O.D. measurements of the overnight cultures to a starting O.D. of 0.0001.

### Monoculture studies

For monocultures, strains were incubated in a 96-well Falcon plate and optical densities were measured every 10 minutes at 600 nm in a Synergy Biotek plate reader heated to 37 °C with continuous shaking. Maximum growth rate and lag time were extracted using a Matlab script written by Daan Kiviet, ETH Zurich (see SI). For statistical analyses, a 3-way ANOVA was conducted (see SI). For the statistical analysis of data in Figure 3, we conducted a 2-way ANOVA for the growth of the recipient (see SI).

### Competition studies

For competition studies, strains were incubated in 24- or 96-well Falcon plates. The initial frequency was set to 1:1 based on O.D. measurements and the initial total O.D. was set to 0.0001. The initial as well as the final ratio of mCherry-to-GFP-tagged strains were measured via flow cytometry. In brief, cells were fixed with 2% formaldehyde, diluted 1:2000, and measured with a Gallios 3-laser flow cytometer (Beckman Coulter, USA). 100’000 events were recorded and cell clusters were computed with an automated clustering algorithm, flow Peaks (Ge and Sealfon, 2012). The clusters were assigned to GFP- and mCherry-tagged cells based on the fluorescence ratios.

### Calculation of relative fitness

Relative fitness was calculated as in (Ross-Gillespie *et al.*, 2007), based on the initial and final frequencies of the strains. Statistical tests were performed with R Version 3.1.2 (R Core Team (2014). R: A language and environment for statistical computing. R Foundation for Statistical Computing, Vienna, Austria. https://www.R-project.org/) with a Wilcoxon-Rank Sum and a Welch t-test (see SI).

### Pyochelin isolation and HPLC analysis

To isolate pyochelin, a slightly modified protocol from (Cox and Graham, 1979) was used. In brief, cells were removed by centrifugation (30 min, 5000 rpm, 4°C) followed by filtration with a 0.2 *μ*m filter. The supernatant was acidified with HCl to pH 1.8-2. Three extractions with 1/3 volume of ethylacetate were performed. The ethylacetate was evaporated with nitrogen, and pyochelin was extracted from the remaining powder by addition of HPLC-grade methanol, rigorous vortexing, and centrifugation of other precipitated substances (5 min, 5000 rpm, room temperature). The supernatant was taken off and directly analyzed on a HPLC (see SI).

### Time course studies

Competition outcome and pyochelin concentration were measured at the same time with one type of strain-color combination (nonrecipient:GFP, secretor:mCherry). Pyochelin was isolated as explained above. The ratios of the strains were estimated by flow cytometry of fixed cells as explained above. At low cell concentrations, fewer events were recorded with the flow cytometer, with a minimum of 5000 events. Also, at some time points the background counts of nonfluorescent cells were high (up to 50% of counts), and the fractions were corrected for this (i.e., we determined the fraction of secretor:mCherry and nonrecipient:GFP among the *fluorescent* cells).

### Thermodynamic speciation calculations

Thermodynamic speciation calculations were performed using the PHREEQC 3 code (Parkhurst and Appelo, 2013) and the Phreeqc thermodynamic database. Stability coefficients and deprotonation constants for pyochelin were taken from (Brandel *et al.*, 2012) and were corrected for zero ionic strength using the Davies equation. All calculations assumed a solution composition of the succinate minimal medium at a constant pH 7.2. Total concentrations of iron and of pyochelin were varied for each analysis. Precipitation of ferrihydrite in equilibrium with the solution was allowed using a solubility coefficient of 10^3.55^ (Schindler *et al.*, 1963).

## Results and Discussion

Our experimental system consisted of three strains based on *P. aeruginosa* and its siderophore pyochelin (Table 1). All strains are knocked out in the production of the primary siderophore pyoverdine, making pyochelin the only siderophore present. The ‘secretor’ strain can secrete and take up pyochelin. The ‘recipient’ does not produce pyochelin, but takes it up. Finally, we constructed the ‘nonrecipient’, which is neither able to produce pyochelin nor to take it up due to a knockout in the pyochelin receptor FptA. We first tested whether the nonrecipient would be inhibited through the addition of pyochelin, in line with the hypothesis that siderophore production could impose growth inhibition in strains that lack the cognate receptor. Indeed, the nonrecipient was affected by the presence of the siderophore (6 *μ*M) in three key growth parameters: Its maximal growth rate and yield were reduced, and its lag phase was prolonged (Fig. 1a, Fig. S2). These effects were linked to the pyochelin receptor knockout, since the recipient was not inhibited by the addition of pyochelin (Fig. 1b). Also, the inhibition was only observed when iron concentrations were limiting (1 *μ*M FeCl_3_; Fig. 1a, Fig. S2).

**Figure 1.**
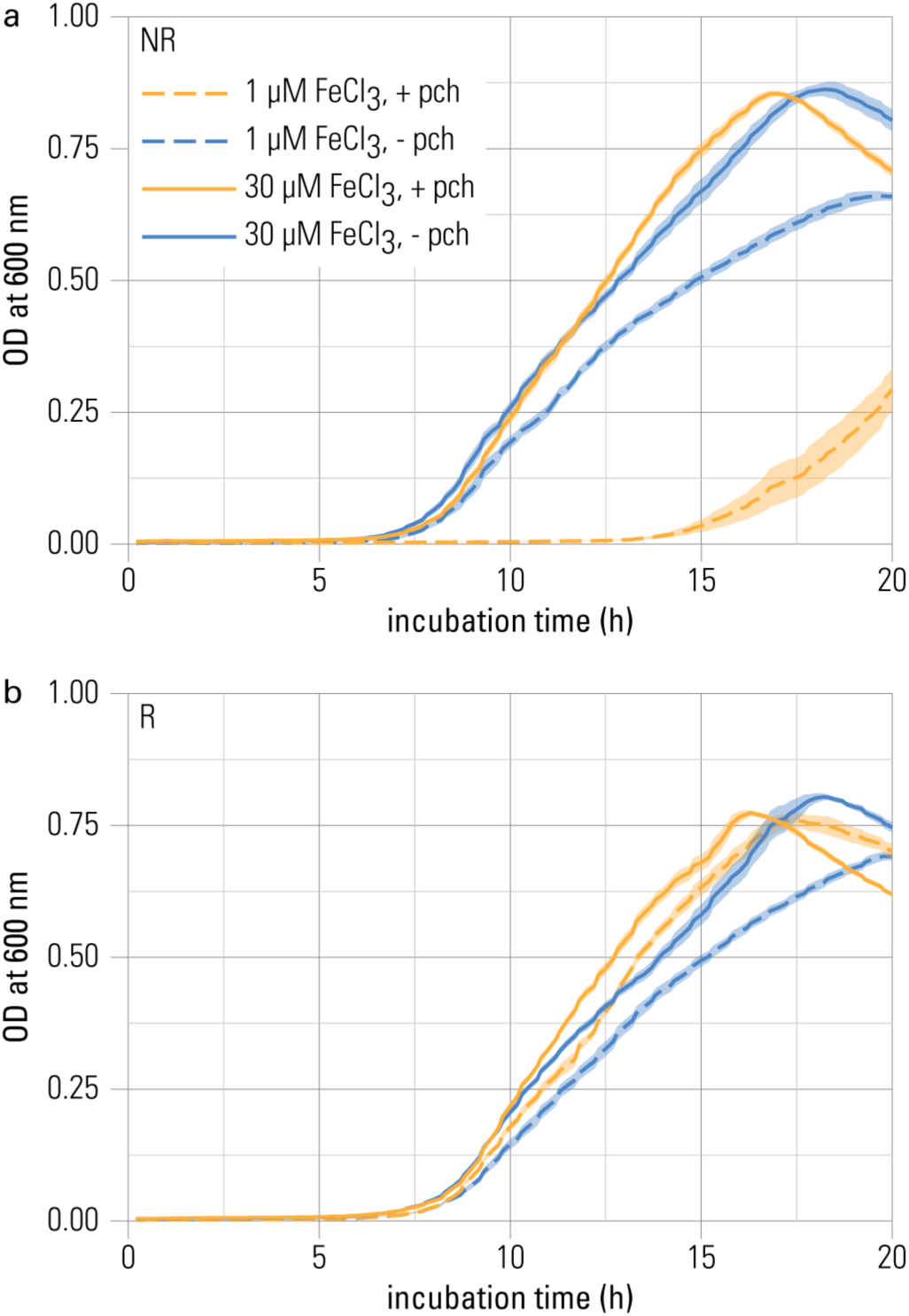
Pyochelin inhibits growth of a strain without pyochelin receptor at iron limitation. (a) To measure the effect of pyochelin on iron competition, three strains were used based on *Pseudomonas aeruginosa* PAO1. All three are knocked out in pyoverdine production, rendering pyochelin the only siderophore in the system. (b) Growth of nonrecipient NR (without pyochelin receptor) is inhibited in the presence of 6 *μ*M pyochelin (+ pch) under iron-limited conditions (orange dashed line). (c) Growth of recipient R (with pyochelin receptor) is not inhibited by 6 *μ*M pyochelin. Data of the control treatment without pyochelin addition (- pch) are shown in blue. Data represents mean optical density (O.D.) measurements of nine replicates and the standard error of the mean indicated as shaded areas around the lines. The growth parameters extracted from these growth curves are depicted in Fig. S1.

**Table 1.** Overview of genetically engineered *Pseudomonas aeruginosa* PAO1 strains used to investigate the competitive effect of pyochelin.

| strain | pyochelin production | pyochelin uptake | relevant knockouts |
| --- | --- | --- | --- |
| secretor (S) | yes | yes | $\Delta pvdD$ |
| recipient (R) | no | yes | $\Delta pvdD\Delta pchEF$ |
| nonrecipient (NR) | no | no | $\Delta pvdD\Delta pchEF$<br>$\Delta fptA$ |

Based on thermodynamic modeling, we propose a mechanism for achieving growth inhibition. The nonrecipient, which is unable to access pyochelin-bound iron, can be inhibited when pyochelin temporarily depletes free aqueous iron species below their equilibrium concentration (Fig. 2). Given enough time, all iron sequestered by pyochelin will be replenished by dissolution of precipitated iron species, and any inhibition of the nonrecipient will be released (see Supplementary Information SI for more detailed discussion). However, if pyochelin binds dissolved iron more rapidly than it is replenished from precipitated phases, and if the uptake rate of iron by the nonrecipient strongly depends on the concentration of dissolved iron (Campbell, 1996), then siderophores can temporarily reduce iron uptake by the nonrecipient and inhibit its growth. Thus, in kinetically controlled, saturated systems siderophores can induce growth inhibition efficiently, even if the siderophore has a low iron-binding affinity or is present at low concentrations (as is the case for pyochelin in our experiments). Kinetically controlled, saturated systems are potentially widespread in the environment, e.g., in marine high-nutrient-low-chlorophyll ocean areas where the iron concentration is controlled by mobilization rates from atmospheric dust inputs (Kraemer *et al.*, 2005).

**Figure 2.**
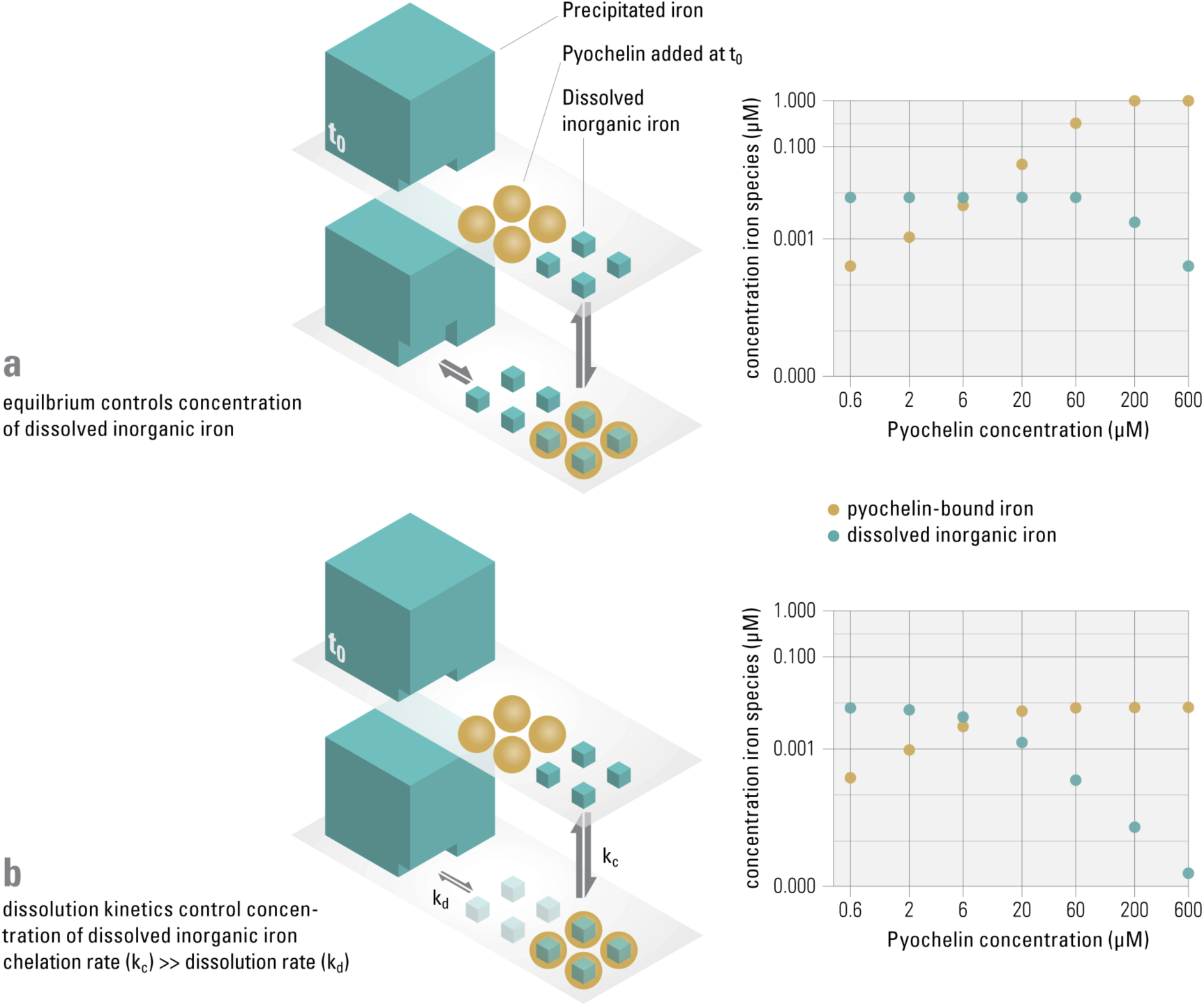
Pyochelin can only sequester dissolved iron sufficiently if the system is not in equilibrium, but controlled by slow dissolution kinetics. In our experimental system, similar to other aerobic, pH neutral environments, 99% of the iron precipitates (i.e., a saturated system), while only around 1% is present as dissolved inorganic iron, which we assume to be available to the nonrecipient (t_0_). We consider the following two extreme scenarios. Equilibrium controlled system (panel a): When pyochelin is added and the system is in equilibrium, the concentration of dissolved inorganic iron is buffered by the precipitated phases and remains constant. Kinetically controlled system (panel b): When iron sequestered by pyochelin is not replenished due to slow dissolution kinetics of the precipitated iron, the concentration of dissolved iron can be significantly reduced. Thermodynamic modeling was used to calculate concentrations of iron species for increasing pyochelin concentrations for both scenarios (data plots on the right). In equilibrium controlled systems, the dissolved inorganic iron concentration stays constant at 0.08 *μ*M in presence of up to 60 *μ*M pyochelin. In dissolution controlled systems, iron sequestered by pyochelin is not replenished on the time-scale of the experiment and the concentration of dissolved iron can be reduced in the presence of 6 *μ*M pyochelin. Experimentally observed growth inhibition of the nonrecipient (Fig. 1) supports a kinetically controlled system. However, if slow dissolution can partially replenish dissolved iron concentrations on the timescales of the experiments, the available iron concentrations to the nonrecipient are expected to be in between these extreme scenarios illustrated (see Supplementary Info for more detailed discussion).

Our hypothesis of kinetically mediated inhibition is consistent with the observation that the nonrecipient started growing after a lag phase of several hours upon addition of pyochelin (Fig. 1a). At longer incubation times, sufficient iron can be released from precipitated phases to support growth or, alternatively, the nonrecipient can adapt to the low iron concentration (e.g., by increasing the number of transporters). Kinetics can also explain the reduced inhibitory effect of pyochelin at a higher iron concentration (30 *μ*M FeCl_3_, Fig 1a, Fig. S2). This response was not caused by higher dissolved inorganic iron concentrations, since the system was already saturated at low iron concentration (Supplementary Table 2). This response is rather a consequence of increased amounts of precipitation that lead to an increased surface area (given that the specific surface area is constant (Dzombak and Morel, 1990) and thus faster iron oxide dissolution (Kraemer, 2004).

After establishing that inhibition can occur in kinetically controlled, saturated systems, our goal was to test whether inhibiting competitors is an important biological function of pyochelin production. To address this hypothesis, we separately assessed the two distinct benefits of pyochelin of (a) making iron more available for growth and (b) making iron unavailable to a competitor. To disentangle costs and benefits, we measured benefits of pyochelin in the recipient, a strain able to acquire but incapable of producing pyochelin and thus paying no costs for its production. We grew the recipient in the presence and absence of pyochelin in both monoculture and in competition with the nonrecipient. We quantified the frequency of the strains in competition by flow cytometry, using strains that constitutively expressed fluorescent markers (P_A1/04/03_::*egfp* or *mcherry*). To assess the benefit of pyochelin, we calculated the final optical density (O.D.) reached by each strain in competition and compared it to the final O.D. reached in monoculture. We found that the addition of pyochelin was only beneficial in competition, since there was no increase in the recipient’s yield in monoculture, but a strong increase in the competition (Fig. 3). These results suggest that in environments where siderophores are not important for increasing iron availability, they might still be very important as a competitive strategy.

**Figure 3.**
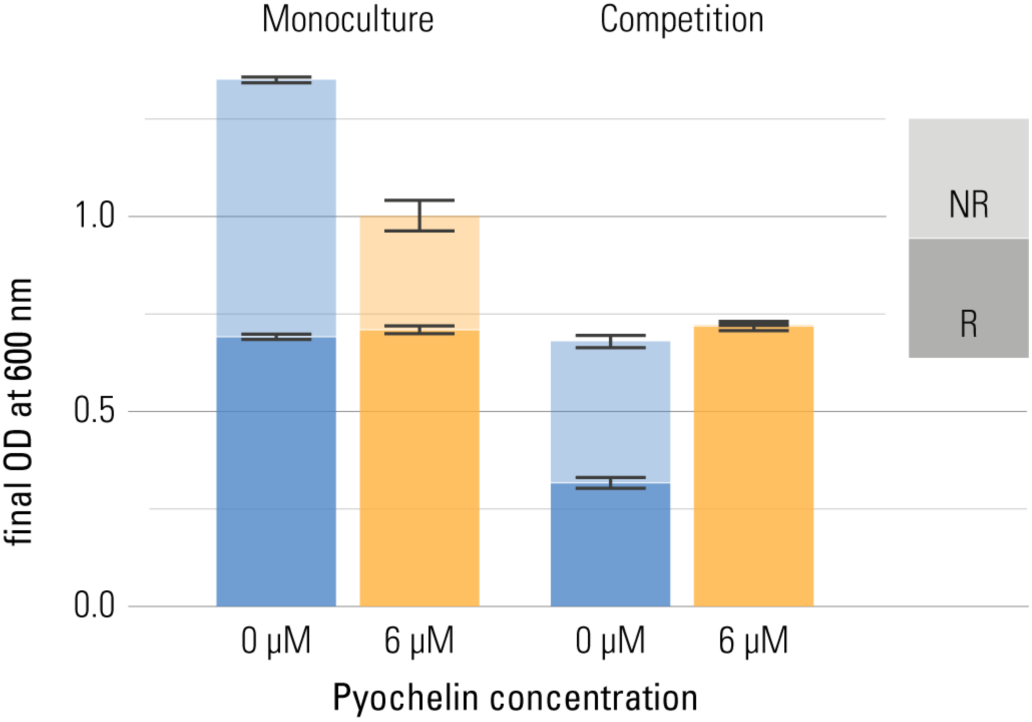
Pyochelin can be highly beneficial in competition, even when it is not beneficial in monoculture. Recipient (R) and nonrecipient (NR) were grown under iron limitation (1 *μ*M FeCl_3_) either in monoculture (left) or in 1:1 competition (right). The effect of pyochelin was tested by addition of 6 *μ*M pyochelin standard (orange; control without pyochelin addition in blue). The final yield of each strain reached after 20 hours of growth, as O.D. at 600 nm, is shown as stacked bars (nine replicates per type; error bars depict the standard error of the mean). We calculated the final O.D. reached by each strain in competition by multiplying the final O.D. of the whole population (recipient+nonrecipient) with the frequency of each strain as determined by flow cytometry. The final yield of the recipient did not significantly increase upon addition of pyochelin in monoculture, but strongly increased in competition (2-way ANOVA shows a significant 2-way interaction between pyochelin addition and social environment for the response variable final O.D., F=442, p < 0.001).

The suppressive effect of pyochelin on the nonrecipient’s growth was aggravated by the presence of the recipient. In monoculture, pyochelin reduced the yield of the nonrecipient only by about a factor of two (from a mean O.D. of 0.67 to 0.29). However, in competition in the presence of pyochelin, the yield of the nonrecipient dropped below the level of detection. Thus, a synergistic effect of pyochelin and a growing recipient strain on the nonrecipient was apparent. Again, this can be explained by a kinetically mediated inhibition effect, given the assumptions we introduced before. Assuming that the recipient recycles pyochelin (as is the case for numerous siderophores (Ratledge and Dover, 2000), recipients would take up the pyochelin-iron complex, remove the iron, and afterwards release unbound pyochelin. This way, growing recipients would constantly create unchelated pyochelin siderophores that bind iron and significantly deplete the dissolved inorganic iron fraction.

In the above-described experiments, pyochelin was added to the medium. To investigate whether bacteria themselves would produce enough pyochelin to suppress competitors, we performed competition studies between a secretor and a nonrecipient with an initial ratio of 1:1. The two strains were again distinguished on the basis of a constitutively expressed fluorescent protein. The fraction of nonrecipient cells decreased after around 12 hours of competition at iron limitation, temporally associated with the onset of pyochelin production (Fig. 4a, b), supporting our hypothesis that growth inhibition is induced by pyochelin sequestering available iron. We additionally quantified the outcome of competition after 20 hours of growth under iron-limited as well as iron-rich conditions (1 *μ*M FeCh_3_ and 30 *μ*M FeCh_3_, respectively) by calculating changes in the relative frequency of the secretor (Fig. 4c). The secretor increased in frequency under low as well as high iron concentration. Control competition experiments performed with the alternative fluorescent protein-strain combination led to the same qualitative outcome. The observed inhibition at high iron concentration shows that siderophores can have a growth-reducing effect even when the environment is not iron limiting *per se*. Therefore, siderophores might mediate competition for resources other than iron, because if a competitor is inhibited in accessing iron, its reduced growth also translates into reduced consumption of other resources.

**Figure 4.**
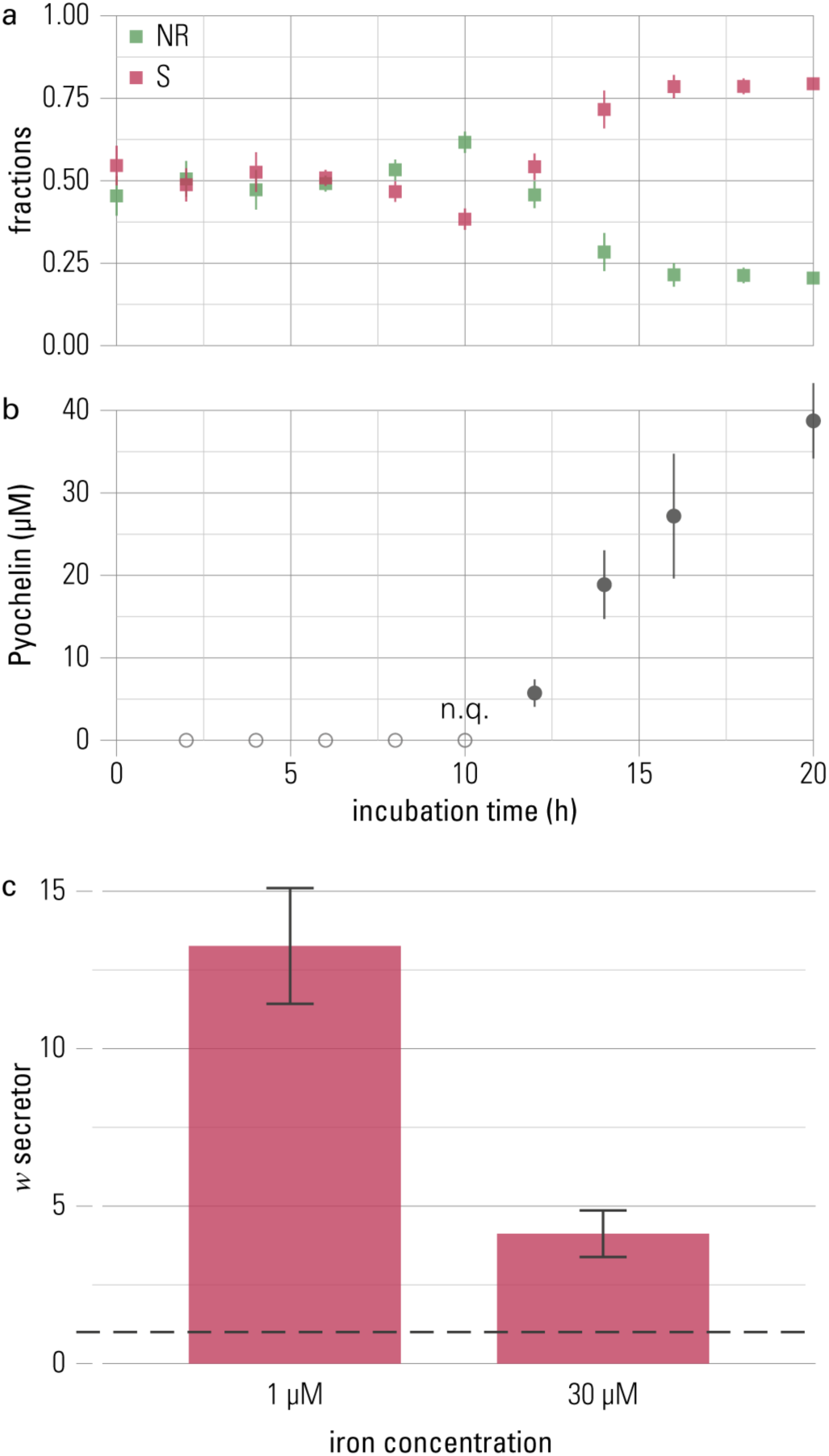
The competitive advantage of the pyochelin-secretor is temporally linked to production of pyochelin. Secretor and nonrecipient were grown in 1:1-competitions for 20 hours. (a) The fractions of each competition partner are shown over time with the secretor (S) in red and the nonrecipient (NR) in green. The frequency of the secretor increased after 12 hours of growth. (b) Increase of the secretor was temporally associated with the production of pyochelin. Pyochelin was detectable in the supernatant after 10 hours and accumulated in the medium over time. Means of three biological replicates are shown and error bars represent standard error of the mean (n.q. = non quantifiable, below quantification limit of 1 *μ*M). (c) To quantify the overall outcome of competition, a proxy for relative fitness (*w*, (Ross-Gillespie *et al.*, 2007)), was calculated. Neutral competition outcome, where *w* equals 1, is indicated by a dashed line, and *w* > 1 indicates higher fitness of the secretor. The secretor shows a higher fitness at both low and high iron concentrations (1 *μ*M FeCl_3_ and 30 *μ*M FeCl_3_, respectively; Welch t-test with alternative hypothesis that true mean of *w* is greater than *μ*=1: for 1 *μ*M FeCl_3_ t=6.67, p<0.001, and for 30 *μ*M FeCl_3_ t=4.23, p=0.002). Under iron-rich conditions, the increase of the secretor is smaller, though, than under iron-limitation (Wilcoxon rank sum test, W=60.5, p=0.003). This outcome is robust to the identity of the combination of strain and fluorescent protein (see Fig. S2). The bar chart shows the mean of eight repeats and error bars represent the standard error of the mean.

## Conclusion

Overall, our findings support the hypothesis that siderophores in environmental systems inhibit other bacteria that lack the specific siderophore receptor system, as supported by previous observational studies (Vachée *et al.*, 1997; Gram *et al.*, 1999; Khare and Tavazoie, 2015). We argue that inhibition is highly efficient in kinetically controlled, saturated systems, and that the competitive benefits obtained by siderophore secretion might surpass the benefit of making iron more available, e.g., if iron is not limiting growth *per se.* Our findings also provide a new hypothesis regarding the presence of exogenous receptor genes in bacteria. Thus far, the evolutionary dynamics in siderophore production and uptake were mainly interpreted from the perspective of the public goods dilemma (Kümmerli and Brown, 2010; Cordero *et al.*, 2012). For example, the observation that strains in the lungs of cystic fibrosis patients maintain siderophore receptors only as long as siderophore producers are present could indicate that strains with receptors are selected for because they cheat on producers and save siderophore production costs (Andersen *et al.*, 2015). Our results suggest a complementary perspective, namely that the presence of siderophore producers forces other strains to maintain receptors to secure access to iron. Expression of a receptor makes siderophore-bound iron accessible, and renders bacteria immune against iron sequestration by competitors.

Our findings underline the importance of siderophore secretion for antagonistic interactions, and show that the same biological trait may be important for cooperative as well as competitive interactions. Detecting genes encoding a given trait is thus not sufficient to draw firm conclusions about the relative importance of cooperative and competitive interactions in microbial communities. Rather, addressing this important question requires functional studies under conditions that are close to the natural situation.

## Author contributions

KS, EMJ, KM and MA designed the research. SMK developed the thermodynamic model. KS and EMJ conducted the experiments. KS and MA conducted the statistical analyses. KS, EMJ, SMK, KM and MA wrote the manuscript.

## Acknowledgements

This work was funded by ETH Zurich, Eawag and a Swiss National Science Foundation grant to M.A. We thank Rolf Kümmerli for providing strains, as well as Nicola Rhyner (Eawag) for integrating the mCherry marker. We also thank Tina Jaeger for advice on the transduction, and Jacob Malone and Urs Jenal for providing the phage. We thank Daan Kiviet for making his growth analysis script available and David Kistler for help with iron measurements. Also, we thank Laura Sigg and Alma Dal Co for helpful discussions and Alejandra Rodriguez for comments on the manuscript, as well as Thierry Sollberger for help with designing the illustrations in Figure 2.

## References

Andersen SB, Marvig RL, Molin S, Krogh Johansen H, Griffin AS. (2015). Long-term social dynamics drive loss of function in pathogenic bacteria. Proc Natl Acad Sci U S A 112: 10756–10761.

Andrews SC, Robinson AK, Rodríguez-Quiñones F. (2003). Bacterial iron homeostasis. FEMS Microbiol Rev 215–237.

Brandel J, Humbert N, Elhabiri M, Schalk IJ, Mislin GLA, Albrecht-Gary A-M. (2012). Pyochelin, a siderophore of Pseudomonas aeruginosa: physicochemical characterization of the iron(III), copper(II) and zinc(II) complexes. Dalton Trans 41: 2820–2834.

Brockhurst MA, Buckling A, Racey D, Gardner A. (2008). Resource supply and the evolution of public-goods cooperation in bacteria. BMC Biology 6: 20.

Campbell PGC. (1996). Interactions Between Trace Metals and Aquatic Organisms: A Critique of the Free Ion Activity Model. In: Metal Speciation and Bioavailability in Aquatic Systems, Tessier, A & Turner, RD (eds) Vol. 3, John Wiley & Sons: New York, pp 45–102.

Cordero OX, Ventouras L-A, Delong EF, Polz MF. (2012). Public good dynamics drive evolution of iron acquisition strategies in natural bacterioplankton populations. Proc Natl Acad Sci U S A 109: 20059–20064.

Cox CD, Graham R. (1979). Isolation of an iron-binding compound from Pseudomonas aeruginosa. J Bacteriol 137: 357–364.

Coyte KZ, Schluter J, Foster KR. (2015). The ecology of the microbiome: Networks, competition, and stability. Science 350: 663–666.

Dzombak DA, Morel FMM. (1990). Surface Complexation Modeling: Hydrous ferric oxide.

Eberl HJ, Collinson S. (2009). A modeling and simulation study of siderophore mediated antagonism in dual-species biofilms. Theor Biol Med Model 6: 30.

Fgaier H, Eberl HJ. (2010). A competition model between Pseudomonas fluorescens and pathogens via iron chelation. J Theor Biol 263: 566–578.

Fgaier H, Eberl HJ. (2011). Antagonistic control of microbial pathogens under iron limitations by siderophore producing bacteria in a chemostat setup. J Theor Biol 273: 103–114.

Ge Y, Sealfon SC. (2012). flowPeaks: a fast unsupervised clustering for flow cytometry data via K-means and density peak finding. Bioinformatics 28: 2052–2058.

Ghysels B, Dieu BTM, Beatson SA, Pirnay J-P, Ochsner UA, Vasil ML, et al. (2004). FpvB, an alternative type I ferripyoverdine receptor of *Pseudomonas aeruginosa*. Microbiology (Reading, Engl) 150: 1671–1680.

Gram L, Melchiorsen J, Spanggaard B, Huber I, Nielsen TF. (1999). Inhibition of Vibrio anguillarum by Pseudomonas fluorescens AH2, a possible probiotic treatment of fish. Appl Environ Microbiol 65: 969–973.

Griffin AS, West SA, Buckling A. (2004). Cooperation and competition in pathogenic bacteria. Nature 430: 1024–1027.

Khare A, Tavazoie S. (2015). Multifactorial Competition and Resistance in a Two-Species Bacterial System Zhang, J (ed). PLoS Genet 11: e1005715.

Kraemer SM. (2004). Iron oxide dissolution and solubility in the presence of siderophores. Aquatic Sciences - Research Across Boundaries 66: 3–18.

Kraemer SM, Butler A, Borer P, Cervini-Silva J. (2005). Siderophores and the Dissolution of Iron-Bearing Minerals in Marine Systems. Rev Mineral Geochem 59: 53–84.

Kümmerli R, Brown SP. (2010). Molecular and regulatory properties of a public good shape the evolution of cooperation. Proc Natl Acad Sci U S A 107: 18921–18926.

Lambertsen L, Sternberg C, Molin S. (2004). Mini-Tn7 transposons for sitespecific tagging of bacteria with fluorescent proteins. Environ Microbiol 6: 726–732.

Oliveira NM, Martinez-Garcia E, Xavier J, Durham WM, Kolter R, Kim W, et al. (2015). Biofilm Formation As a Response to Ecological Competition Laub, MT (ed). PLoS Biol 13: e1002191.

Parkhurst DL, Appelo CAJ. (2013). Description of input and examples for PHREEQC version 3: a computer program for speciation, batch-reaction, one-dimensional transport, and inverse geochemical calculations. In: Techniques and Methods, p 497.

Ratledge C, Dover LG. (2000). Iron metabolism in pathogenic bacteria. Annu Rev Microbiol 54: 881–941.

Ross Gillespie A, Gardner A, West SA, Griffin AS. (2007). Frequency Dependence and Cooperation: Theory and a Test with Bacteria. Am Nat 170: 331–342.

Ross-Gillespie A, Gardner A, West SA, Griffin AS. (2007). Frequency Dependence and Cooperation: Theory and a Test with Bacteria. Am Nat 170: 331–342.

Schindler P, Michaelis W, Feitknecht W. (1963). Loeslichkeitsprodukte von Metalloxiden und-hydroxiden. 8. Mitteilung. Die Loeslichkeit gealterter Eisen(III)-hydroxid-Faellungen. Helv Chim Acta 46: 444–449.

Simões M, Simões LC, Pereira MO, Vieira MJ. (2008). Antagonism between Bacillus cereus and Pseudomonas fluorescens in planktonic systems and in biofilms. Biofouling 24: 339–349.

Traxler MF, Seyedsayamdost MR, Clardy J, Kolter R. (2012). Interspecies modulation of bacterial development through iron competition and siderophore piracy. Mol Microbiol 86: 628–644.

Trejo Hernandez A, Andrade-Dominguez A, Hernandez M, Encarnacion S. (2014). Interspecies competition triggers virulence and mutability in Candida albicans–Pseudomonas aeruginosa mixed biofilms. ISME J 8: 1974–1988.

Vachée A, Mossel DA, Leclerc H. (1997). Antimicrobial activity among Pseudomonas and related strains of mineral water origin. J Appl Microbiol83: 652–658.

Velicer G. (2003). Social strife in the microbial world. Trends Microbiol 11: 330–337.

Wandersman C, Delepelaire P. (2004). Bacterial Iron Sources: From Siderophores to Hemophores. Annu Rev Microbiol 58: 611–647.

West SA, Griffin AS, Gardner A, Diggle SP. (2006). Social evolution theory for microorganisms. Nature Rev Microbio 4: 597–607.

